# Expectation management in humans and LLMs

**DOI:** 10.1101/2025.03.09.642185

**Authors:** Benjamin Menashe, Austin Drake, Michal Ben-Shachar

## Abstract

Mirative markers, such as “surprisingly”, explicitly encode a violation of expectations. Such markers are used for expectation management during communication. Sensitivity to mirative markers relies on two abilities: i) updating expectations upon recognizing a mirative marker, and ii) identifying expectation violations warranting the use of a mirative marker. In this study, we compared sensitivity to mirative markers in humans and large language models (LLMs). In part 1, we used a sentence-completion task, where humans and LLMs were presented with sentence fragments and asked to continue them. Results show that for both humans and LLMs, the presence of a mirative marker significantly increased response entropy and decreased top-response probability, in line with theoretical accounts of mirative processing. In part 2, we created a novel task, mirative polarity selection, where humans and LLMs are presented with a sentence pair and asked to select whether it was connected by a mirative marker (“surprisingly”) or an anti-mirative marker (“unsurprisingly”). Results show that LLMs perform at an impressive human-equivalent level. We conclude that both humans and LLMs use mirative markers as cues for calibrating their subsequent expectations during sentence processing.

## Introduction

Humans constantly form expectations about upcoming stimuli^1,2^. Linguistic stimuli are no exception: it is well-demonstrated that humans use context and prior knowledge to form expectations and predictions about upcoming linguistic information, and then test those predictions against the actual upcoming input^3–5^. Encountering a violation of semantic expectations is associated with longer processing time^6–10^, an elevated amplitude of the N400 event-related potential (ERP) component^11–14^, and neural activation across many brain regions throughout the language network^15–19^. These results are in line with “predictive processing” theories of the brain, whereby such violations of expectations are regarded as prediction errors which the system attempts to minimize by calibrating expectations^20–23^.

In natural discourse, speakers often attempt to manage their interlocutors’ expectations explicitly, using *mirative markers*. Mirative markers are linguistic elements that encode violation of expectations^24–26^, including words such as “surprisingly” and “remarkably”. Mirative markers are widely used to inform about an upcoming unexpected message. In theory, such expectation management may help interlocutors avoid (or reduce) prediction errors and the associated increase in processing time and difficulty incurred by such errors.

In recent years, large language models (LLMs) have revolutionized the field of natural language processing. LLMs provide a probability estimate, for each token in their vocabulary, reflecting that likelihood that this token would follow a given sequence of text. Such predictions are based on exposure to enormous corpora of language, during training^39–42^. This process has been compared to forming expectations in humans, in what has been called a “shared computational principle of language comprehension”^43^. LLMs have been previously used to gain insight into human behavior^44,45^ and neural activity^46–48^.

This study investigates expectation management in humans and LLMs, by quantifying and comparing their sensitivity to mirative markers. Sensitivity to mirative markers relies on two abilities: i) updating expectations upon recognizing a mirative marker, and ii) identifying expectation violations warranting the use of a mirative marker. Comparing humans and LLMs serves two purposes: first, if LLMs demonstrate a similar sensitivity to mirative markers as humans, LLMs could serve as efficient “test subjects” for future explorations of expectation management. Second, expectation management is theorized to be a byproduct of predictive processing, as it is hypothesized to reduce prediction error. Since humans and LLMs both form predictions during discourse (while being otherwise radically different), observing emergent expectation management in LLMs would provide evidence that it is a byproduct of predictive communication systems.

*Mirativity* is considered to be a language-universal category^24^. It can be marked using words (e.g., “surprisingly”), exclamatory particles or intonation (e.g., “Wow!”), or, in some languages, by specialized morphological affixes^27^. Conversely, a confirmation of expectations can also be marked using words such as “unsurprisingly” and “expectedly”, which we refer to as *anti-mirative markers*. According to linguistic theory, mirative markers are licensed only when there exists a more expected proposition that is false^28^. This implies that mirative markers modify expectations by decreasing the probability of the most-expected upcoming stimuli and increasing the probability of lesser-expected upcoming stimuli. Such a process is referred to as *domain widening*, whereby the domain of plausible upcoming stimuli is expanded to include lesser-expected stimuli^29,30^. We propose that this is effectively equivalent to an increase in the entropy of the distribution of expectations (a quantitative measure of uncertainty^31^). The effects of increased entropy on human behavior and neural activity have been studied extensively^32–36^.

Two previous studies that addressed the neural effects of expectation-disconfirming markers, both using ERPs, report conflicting results. In the first study^37^, participants read scenarios containing unexpected target words, which were preceded by the discourse marker “even so” on half the trials. Results showed that the presence of “even-so” attenuated the amplitude of the N400 component for otherwise unexpected target words. In a later study^38^, participants read dialogues in Dutch with unexpected endings, which were preceded by the discourse marker “eigenlijk” (“actually”) on half the trials. Results showed no evidence for a main effect of discourse marker on the N400 amplitude, nor an interaction effect of the discourse marker and predictability on the N400 amplitude. As such, there is currently only limited and conflicting empirical evidence on processing mirative markers and subsequent expectation modification.

In part 1 of this paper, we studied how mirative markers modify expectations. We used a sentence-completion task, where human participants and LLMs were presented with sentence fragments and asked to continue them. We manipulated the sentences to begin either with a mirative marker or a neutral marker, and compared the respective responses. Since a mirative marker indicates an upcoming violation of expectations, we predicted that it would lead to a decrease in the probability of the most-expected response and to a general increase in the probability of lesser-expected responses, leading to an overall increase in the entropy of responses, for both humans and LLMs. In part 2, we studied whether LLMs identify violations of expectations to the same level as human participants do. We created a novel task, *mirative polarity selection*, where human participants and LLMs are presented with a sentence pair and asked to select whether it was connected by a mirative marker or by an anti-mirative marker. We predicted humans would perform well on the task, and that LLMs would perform near or at a human-equivalent level, in line with the performance of LLMs on theory-of-mind tasks^51^.

Altogether, the two tasks allowed us to examine the core conditions required for successful expectation management using mirative markers: their ability to modify subsequent expectations appropriately and the capacity to deploy them in appropriate contexts.

## Results

### Part 1: Increased entropy in response to mirative markers

Using a sentence-completion task, we tested how the presence of a mirative marker (“surprisingly”) at the beginning of a sentence fragment, compared to a neutral marker (“yesterday”), affects expectations for upcoming words. We tested human participants using an online questionnaire that contained thirty novel sentence fragments. We then tested the Meta-Llama-3/3.1/3.2 series of LLMs on the same sentence fragments.

In human participants, we found that the inclusion of a mirative marker in the beginning of a sentence fragment resulted in a significant *decrease* in the probability of the most common response to the neutral marker (t_28_ = −6.85, p<.001, Fig. 1A), and a significant *increase* in the entropy of the response distributions (t_28_ = 5.58, p<.001, Fig. 1B. See also Fig. 1C for an example of response distributions). In LLMs, similarly to human participants, mirative markers evoked a significant decrease in probability of the top response (see Fig. 1D for an example LLM and Supplementary Table S1 for the full statistics of all LLMs tested). Moreover, the entropy of the response distribution following a mirative marker increased for all LLMs, although this increase was significant in only three of the six LLMs tested (Llama-3-8B, Llama-3.1-8B, Llama-3.2-3B; See Fig. 1E for an example LLM). A visual inspection of the top 10K tokens suggested that mirative markers decreased the probability of the top 10-20 tokens and increased the probability of the rest of the tokens (Fig. 1F). Taken together, this provides evidence that humans and LLMs process the mirative marker as an indicator for an upcoming violation of expectation, and both humans and LLMs modify their expectations accordingly.

**Figure 1.**
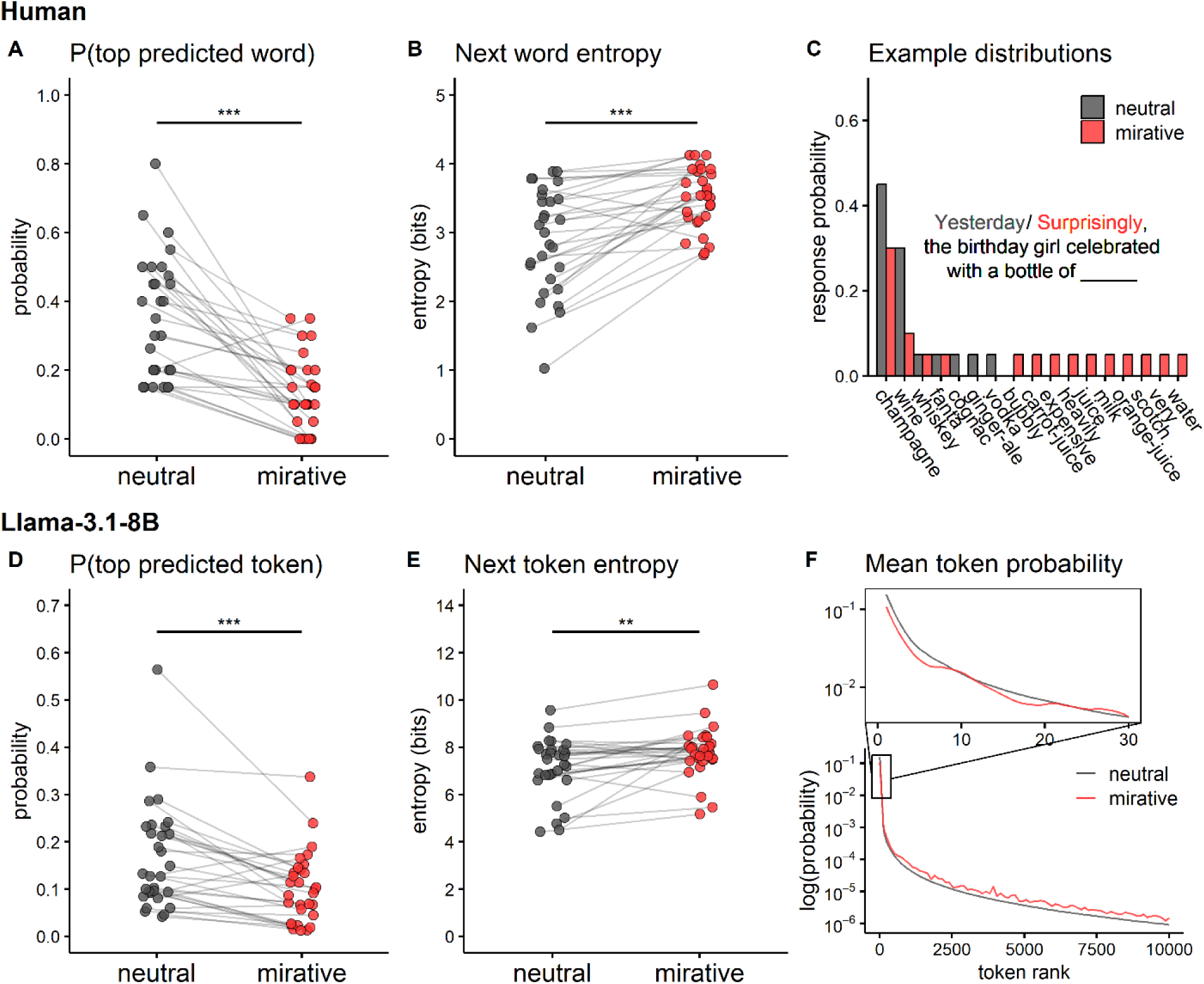
Mirative markers modify expectations. Top: Human participants were presented with 30 sentence fragments and asked to continue them with the next word. Stimuli began either with a neutral marker (“yesterday”) or a mirative marker (“surprisingly”). **A)** Probability of the most common response in the neutral condition, and of the same response in the mirative condition. **B)** Entropy over the response distribution, per condition. **C)** Example of response distributions for the same sentence fragment (“the birthday girl celebrated with a bottle of”) in the neutral condition in gray and in the mirative condition in red. **Bottom:** We used an LLM (Meta-Llama-3.1-8b) to generate next token predictions for the same 30 sentence fragments presented to human participants. **D)** Probability of the top token in the neutral condition, and the same token in the mirative condition. The difference is significant even when removing the top outlier point. **E)** Entropy over the next token distribution, per condition. **F)** Mean probability by marker, calculated across all stimuli, for each of the top 10K tokens in the neutral condition. X-axis represents token rank, determined by descending probability in the neutral condition. Y-axis is on a log scale. Data points were smoothed using locally estimated scatterplot smoothing (LOESS) with span=10. The upper panel is a magnified view of the top 30 ranks. *** p<.001 ** p<.005.

In a follow-up analysis, we generated 1,365 text fragments of varying lengths taken from the Provo corpus^49^. We defined a “window of altered expectations” as the location and range of the increased entropy due to the presence of a mirative marker. The distance from the mirative marker was measured by number of sentences using a linear mixed-effects regression, and separately by number of words using a spline regression. Results from both methods converge to show that the window of altered expectations extends up to 2 sentences ahead (Fig. 2A) and approximately 5-40 words ahead (Fig. 2B, see also Supplementary Table S2 for the full statistics). This suggests that mirative markers have a limited window in which the violation of expectations is anticipated to occur.

**Figure 2.**
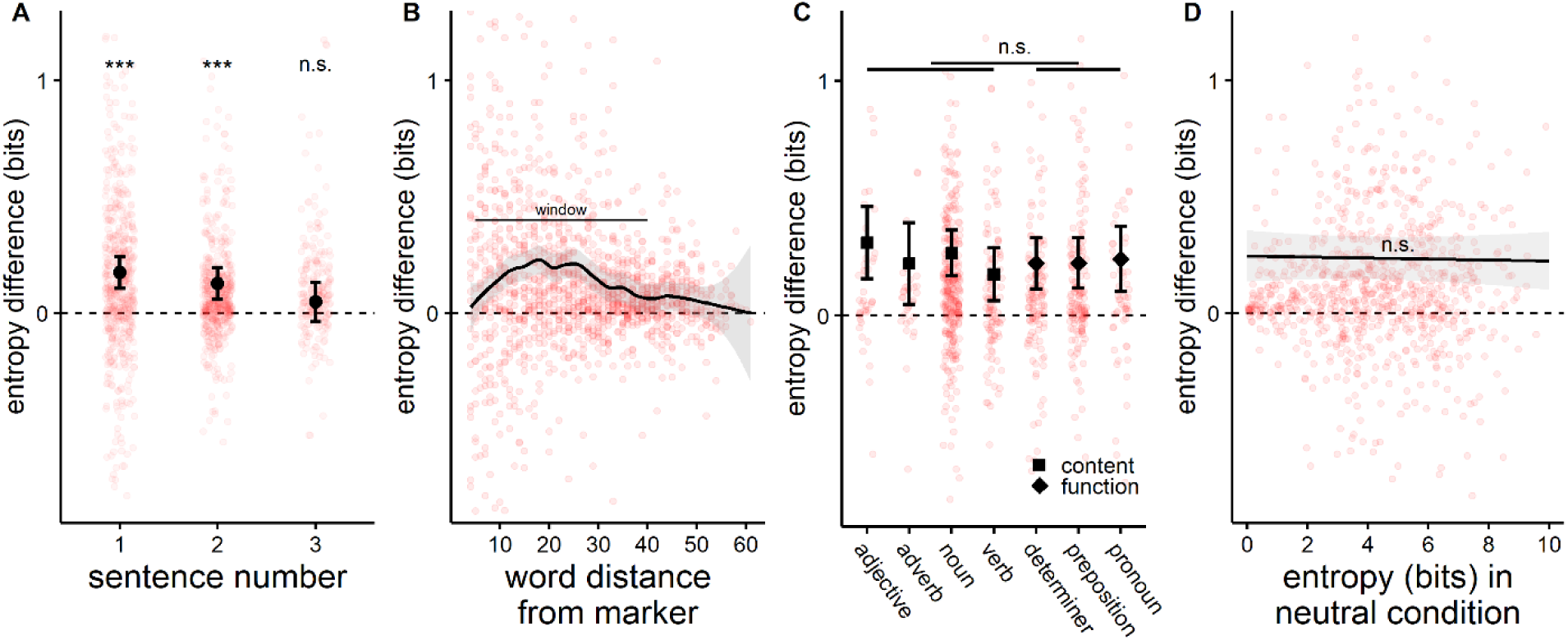
Mirative markers increase entropy. We used an LLM (Meta-Llama-3.1-8B) to quantify the entropy following each word in 1,365 text fragments taken from the Provo corpus (Luke & Christianson, 2018). Texts began either neutrally, i.e., as they were in the corpus, or with a mirative marker (“surprisingly”). All plots show the difference in entropy between the mirative and the neutral condition, per word. **A)** Entropy difference per sentence number. Black dots show the estimated means from a linear mixed-effects model. Error bars show the 95% confidence interval (CI). Outliers above (n=51) and below (n=38) the y-axis range are not shown but were included in the model. **B)** Entropy difference by the word’s distance from the mirative marker. Black line shows the locally estimated scatterplot smoothing (LOESS) with span=20 over the entropy difference of all words. The gray surface around the black line shows the 95% CI. Outliers above (n=51) and below (n=38) the y-axis range are not shown but were included in the locally estimated scatterplot smoothing model. **C)** Entropy difference for each predicted token, categorized by part-of-speech. Black icons show the estimated means from a linear mixed-effects model fit separately for each part-of-speech, with squares for content words and diamonds for function words. Error bars show the 95% CI. Outliers above (n=26) and below (n=17) the y-axis range are not shown but were included in the model. **D)** Entropy difference by the entropy in the neutral condition. Black line shows the estimated trendline from a linear mixed-effects model. The gray surface around the black line shows the 95% CI. Outliers above (n=26) and below (n=17) the y-axis range are not shown but were included in the model. *** p<.001, n.s. not significant.

In a linear mixed-effects model predicting the increased entropy by the POS of the most-expected word, all POS showed a significant increase in entropy and no significant difference was found between content words and function words (t_830_ = 0.38, p=.705, Fig. 2C). Additionally, we wanted to check if mirative markers modify expectations differently for different levels of contextual constraint, measured by the entropy in the neutral condition. No correlation was found between the level of contextual constraint and the increase in entropy in the mirative condition (standardized correlation coefficient = −0.010, p=.774, Fig. 2D). This suggests that the addition of a mirative marker generally results in an increase in entropy, regardless of the level of contextual constraint. Importantly, this also shows that the increase in entropy is not due to regression to the mean.

### Part 2: Identifying expectation violations

We tested human and LLM performance in identifying violations of expectation. For that, we devised a novel “mirative polarity selection” task. Participants and LLMs were presented with a sentence pair and asked to select whether it was connected with a mirative marker (“surprisingly”) or an anti-mirative marker (“unsurprisingly”). Sentence pairs were sampled from the Discovery dataset^50^, and were originally connected by one of the markers in the corpus (this constitutes the correct response). We tested human participants using an online questionnaire, and we used the Meta-Llama-3/3.1/3.2-instruct series and the chatGPT-3.5/4/4o series of LLMs.

Human mean accuracy on the mirative polarity selection task was 75.5% ± 1%. Participant performance varied widely (Fig. 3B), ranging between 46.7% to 90%. This suggests high variability in the participants’ ability to identify violations of expectations. Participant agreement on the responses was fair (percent agreement: 68.2%, Gwet’s AC1: 0.376, significantly better than chance agreement, p<.001). The relatively low level of mean accuracy and agreement may reflect the natural stimuli we used, as the sentence pairs may not have contained sufficient information for selecting the correct marker (see Supplementary Table S5 for examples with low human accuracy). Thus, we take the mean human accuracy and the top human accuracy as tentative benchmarks for this task.

**Figure 3.**
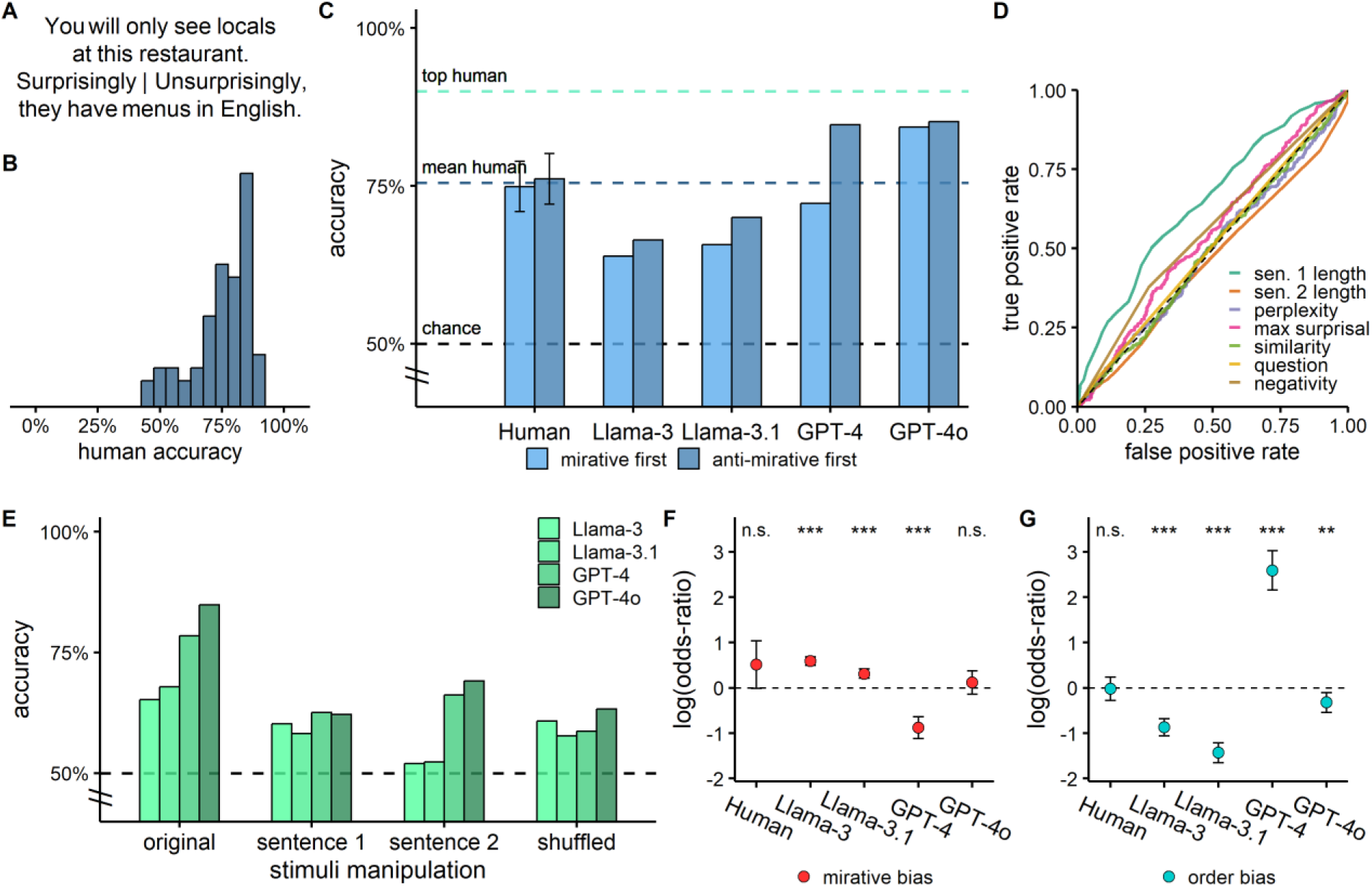
LLMs achieve near human performance on mirative-polarity selection. Mirative-polarity selection is a novel task where participants and LLMs are presented with a sentence pair and asked to select whether it was connected with a mirative marker (“surprisingly”) or an anti-mirative marker (“unsurprisingly”). We used 1000 sentence pairs (500 per marker) from the Discovery dataset (Sileo et al., 2019). Human participants were presented with 60 randomly selected stimuli (30 per marker). **A)** Example stimulus, presented in a mirative-first order. The correct choice is the mirative marker (“surprisingly, they have menus in English”). **B)** Distribution of human accuracy levels in the mirative-polarity selection task. **C)** Accuracy levels achieved by human participants and several LLMs. Choice order with the mirative marker presented first is shown in light blue, and the reverse choice order, with the anti-mirative marker presented first, is shown in dark blue. Error bars indicate a 95% CI for human performance levels. **D)** Area under the curve (AUC) for several discrimination criteria are all close to chance level. Criteria include: length of first or second sentence in tokens, the perplexity and the max surprisal of the second sentence, the cosine similarity between the sentences, whether the first sentence is a question, and the presence of negation in any of the sentences. **E)** LLM mean performance on several conditions of manipulated stimuli. Sentence 1: LLMs were presented with the first sentence only and asked to select which marker, mirative or anti-mirative, is most likely to follow it. Sentence 2: similar manipulation, presenting only the second sentence and asking which marker is likely to precede it. Shuffled: LLMs were presented with random combinations of first and second sentences of different pairs. **F-G)** Mirative bias and choice-order bias estimates, per model, obtained using logistic mixed-effects models. **F)** Mirative bias: positive values suggest a higher probability of selecting the mirative marker over the anti-mirative marker, and negative values vice versa. **G)** Order bias: positive values suggest a higher probability of selecting the second choice presented over the first, and negative values vice versa. Error bars show the 95% CI. ^n.s.^ not significant. ** p<0.01. *** p<.001.

LLM task performance varied, with some models approaching and some exceeding mean human performance (see Supplementary Table S3 for the full statistics, and Fig. 3C for a comparison between human performance and selected LLMs). Within the Meta-Llama series, Llama-3-8B-Instruct and Llama-3.1-8B-Instruct performed well above chance but still below human level. The newer Llama-3.2 models both performed at a chance level. This result may be due to their smaller number of parameters. Within the chatGPT series, GPT-3.5 performed above chance level but below human level, GPT-4 performed at human level, and GPT-4o performed above the mean human level and near the top human level. GPT-4o-mini saw a drop in performance, which may also be due to its smaller number of parameters (see Table S3). A stricter measure of accuracy is the joint order accuracy, where LLM responses were considered correct only if they selected the correct label twice for a given sentence pair (once for each choice order presented). LLM performance as measured by joint order accuracy was lower than model accuracy for all models, although many models still performed above chance level and GPT-4o still performed at an impressive mean human level (see Supplementary Table S3 for the full statistics).

To examine whether the high accuracy of LLMs reflects their ability to identify expectation violations or some other variable, we measured the discriminability of the sentence pairs using several simple variables. Discriminability, measured using area under the curve (AUC), was very poor for most of the variables checked (second sentence length: 0.462, perplexity: 0.492, max surprisal: 0.546, similarity: 0.495, question: 0.508, negativity: 0.558). One exception was the length of the first sentence: anti-mirative markers tended to be preceded by longer sentences as compared to mirative markers (AUC = 0.647, see Fig. 3D for the ROC of the various criteria). However, discriminability based on the length of the first sentence is still not sufficient to explain the performance of the better performing LLMs, as the classification accuracy at the optimal threshold was only 61.4%. Moreover, the length of the first sentence was not a significant predictor of the LLM response for the errors the LLMs made (using a logistic regression with a covariate for stimulus label, p>.42 for all LLMs). This supports the claim that LLMs succeed in mirative polarity selection due to some form of sensitivity to mirativity.

To further assess this claim, we tested LLM performance on manipulated versions of the stimuli, presenting just the first or just the second sentence of the pair. LLM performance decreased substantially in both conditions (Fig. 3E, see Supplementary Table S4 for the full statistics). This also suggests that LLM performance is not due to data contamination (where the model was trained on the stimuli and thus was “supervised” for the correct label), as even one sentence is unique enough to allow retrieval of the appropriate connector if the testing data was memorized. Next, we presented LLMs with sentence pairs where the first and second sentence were randomly shuffled between pairs within the same label (mirative or anti-mirative). The goal of this condition was to eliminate the semantic relation between sentences within each pair, while preserving other global properties (e.g., possible syntactic structure, common function words, register, sentiment…) which might differ between labels. LLM performance decreased substantially on this condition as well (Fig. 3E, see Supplementary Table S4 for the full statistics). This shows that LLM performance is tied to the semantic information in the sentence pair, most likely the occurrence of an expectation violation (or confirmation). Altogether, results from the manipulated stimulus conditions support the view that LLMs succeed in the original task due to some form of sensitivity to mirativity, not due to data contamination or sensitivity to other properties of the stimuli.

*Mirative bias:* Participants tended to select the mirative marker slightly more than the anti-mirative marker, although this difference was not significant (p=.055). Importantly, the mirative marker “surprisingly” is an order of magnitude more frequent than the anti-mirative marker “unsurprisingly” in natural corpora (the Zipf frequency of “surprisingly” is 4.09, compared to 3.01 for “unsurprisingly”, resulting in a frequency ratio of 12.02 to one; these estimates were retrieved using the *wordfreq* python library^52^). This suggests that some mirative bias may be expected if the base frequency of the markers is taken into account when selecting the markers. Of the LLMs which performed above chance level, different tendencies were observed: Llama-3-8B-Instruct and Llama-3.1-8B-Instruct displayed a significant bias towards selecting the mirative marker (p<.001), GPT-4o was not significantly biased towards any label (p=.378), and GPT-4 was significantly biased to select the anti-mirative marker (p<.001, Fig. 3F, see Supplementary Table S3 for the coefficient values). Overall, the different bias tendencies suggest that while some LLMs perform at or above human level, they likely do so through different processes than humans (and of other LLMs).

*Order bias:* LLMs also differed from human participants in their sensitivity to the order of the choices presented (Fig. 3G, see Supplementary Table S3 for the coefficient values). In humans, no significant effect of order was found when estimating using all human participants (N=63; coefficient =-0.02 ± 0.1, p=.880) or when only using the subset of participants (N=22) who were presented with both orders (coefficient = 0.10 ± 0.2, p=.513). In contrast, all LLMs displayed a significant order bias: Llama-3-8B-Instruct and Llama-3.1-8B-Instruct were more likely to select the first choice presented (p<.001), GPT-4o was also more likely to select the first choice presented albeit to a lesser degree (p=.004), and GPT-4 was more likely to select the last choice presented (p<.001). This is consistent with previous reports of order biases in LLMs^53,54^. The fact that LLMs, but not humans, display an order bias in this task further supports the idea that LLMs do not attain their high accuracy in a similar way to humans.

### General discussion

In this study, we assessed sensitivity to mirativity in both human participants and LLMs. We measured how the presence of a mirative marker (“surprisingly”) affects subsequent expectations, and whether humans and LLMs are able to detect expectation violations warranting the use of a mirative marker.

Results from a sentence-completion task show that both humans and LLMs use mirative markers as an indication of an upcoming expectation violation. Results from the mirative-polarity selection task show that both humans and the best-performing LLMs display a similar level of sensitivity to mirativity.

Within “predictive processing” theories^20,22^, there is a clear advantage for giving a warning signal prior to an expectation violation, which allows a preemptive modification of expectations, thus reducing the would-be prediction error. Within linguistic pragmatic theory, Grice’s cooperative principle of communication asserts that speakers aim to help their interlocutors process the conveyed message^55^.

Along these lines, a speaker may use a mirative marker as part of the cooperative principle to help their interlocutor reduce their prediction error for an upcoming unexpected message. Thus, sensitivity to mirativity can provide an advantage for more efficient processing of information. Further studies can directly test this claim by measuring the effect of mirative markers on behavioral and neural correlates of prediction error.

As shown in part 1, human participants modified expectations upon encountering a mirative marker, as did LLMs. The expectation modification involved a reduction (but not an elimination) in the probability of the most-expected response, and an increase in entropy. The reduction in probability of the most-expected response is easily explained, as the literal meaning of mirative markers such as “surprisingly” is that the current message is not the most-expected one^30,28^. The increase in entropy can be interpreted as a general increase in the uncertainty as to the identity of the next upcoming word. This is because, while mirative markers signal an upcoming expectation violation, they do not specify the content of the violation, leading the interlocutor to increase their expectations for many different upcoming words, all of which violate expectations. Additionally, mirative markers do not specify the exact location of the upcoming expectation violation (only that it will appear soon). During word-by-word prediction, this prevents the interlocutor from discounting highly expected words, since the violation could appear later on. Instead, it leads to *domain widening*, where the set of plausible next words expands while preserving the most-expected words^29,30^. Domain widening is congruent with the increase in entropy observed. Moreover, when comparing the top tokens of the LLM responses, overall token rankings remained stable but the probability decreased for the top 10-20 tokens and increased for the rest of the tokens. This supports the notion that uncertainty about the location and identity of the expectation violation leads to expanded expectations when predicting the next word.

In a follow-up analysis, we investigated factors that influence the change in expectations following a mirative marker using a large number of sentence fragments, in LLMs only. Using language models for measuring predictability and entropy of expectations is a standard method that can produce hypotheses regarding the human performance^56,45,43^. Results show a “window of altered expectations” (where mirative markers led to increased entropy in next word predictions) extending approximately two sentences or 5-40 words ahead. Notably, the stimuli were not designed to contain expectation violations, as the Provo corpus passages were taken from a variety of sources (e.g., news articles, popular science magazines) designed for naturalistic reading^49^, to which we added, artificially, a preceding mirative marker. We theorize that, in natural settings, mirative markers may alter expectations until encountering a violation, which is likely to occur within two sentences. The entropy increase was of similar magnitude for expected content words and function words. For example, for the sentence fragment “All that the brain has to work with are”, the most-expected token was “the”, which is a function word (a determiner). When preceded by the mirative marker “surprisingly”, entropy nonetheless increased, although the word “the” is not related to the semantic content of the message. This result may be because less-expected content words remain possible in such positions (for example, “neurons”), or because mirative markers create a general uncertainty in expectations. Since our human experiments were designed to elicit nouns, we are not able to assess conclusively the effect of word type (content or function word) and POS in human participants. Furthermore, entropy increase was independent of the availability of semantically constraining context. The fact that even lower-constraining sentence fragments produced an increase in entropy when preceded by a mirative marker suggests that mirative markers alter expectations at all levels, even weakly held ones. Additionally, this affirms that the increase in entropy is not due to regression to the mean (where extreme values of entropy would have become more moderate).

As shown in part 2, LLMs reached human performance level in selecting the mirative marker as the appropriate connector between a pair of sentences. Identifying if a sentence pair contains a violation of expectations is complex and requires several cognitive abilities: reading comprehension, detecting (often implicit) expectations by applying world knowledge and theory-of-mind in context, and recognizing mismatches between the expectations and the actual message. Thus, success on the task is far from trivial. Human performance varied considerably, ranging from chance level to 90% accuracy.

This could reflect individual differences in the ability to detect expectation violations. It could also reflect differences in the specific expectations people form due to different life experiences, which would lead some to judge that a certain message contains a violation of expectations while others would disagree (but notice that the inter-subject agreement was around 68%, significantly above chance). It could also reflect the possibility that human participants differed in the level of attention and task engagement.

LLMs performed impressively, reaching and exceeding average human level (although still below top human performers). A possible concern is that the stimuli used, taken from the internet, could have been in the training data for the LLMs we tested (“data contamination”). This is unlikely for several reasons: (a) LLM performance is at human-average rather than at ceiling level, (b) LLM’s accuracy dropped when tested on the first or second sentence only (which are sufficient by their own for retrieving a memorized sequence), and (c) the order of the options influenced LLM selection systematically, which is unexpected if the LLM is operating based on memorized examples. An AUC analysis did not reveal any superficial variable that could be used to perform the task (e.g., sentence length, see Fig. 3D). Moreover, performance dropped when sentence pairs were shuffled, ruling out some non-semantic property of the stimuli that the LLMs could have used to classify them. Additionally, a supplementary analysis revealed that the presence of a mirative marker affected LLMs’ responses in a common-sense reasoning task where detecting expectation violations is involved, but not in a vocabulary task where detecting expectation violations is irrelevant for task performance (see Supplementary Fig. S1 in Supplementary Materials). Taken together, this evidence suggests that the success of LLMs in performing the mirative polarity selection task is likely due to their ability to identify expectation violations, which is a core part of sensitivity to mirativity.

The astounding breakthrough success of LLMs in recent years has raised questions about their ability to construct a world model^57,58^ and a theory-of-mind^59,51,60^. A world model can be defined as an internal representation of the properties and causal dynamics of the physical world. Theory-of-mind can be defined as a model of the mental states of another agent, including their beliefs and expectations.

Success in the mirative polarity selection task requires a basic level of both a world model and a theory-of-mind, because, in order to identify what would constitute an expectation violation, one must first have a good approximation of the expectation formed by their interlocutor. There is ample evidence showing that humans possess an understanding of “folk physics”, even at a young age^61–63^. Thus, a world model is helpful in informing the speaker about what their interlocutor expects to happen, under normal circumstances. In an example stimulus, such as: “I saw the issue. Surprisingly/Unsurprisingly, it was easy to fix”, to correctly select the mirative marker (“surprisingly”), one must first have a world model wherein issues are usually difficult to fix, and second, use their theory-of-mind to understand that their interlocutor would also expect the issue to be difficult to fix. The success of LLMs on the task suggests that LLMs may have some sort of a functioning theory-of-mind and informative world model (albeit not necessarily similar in representation to humans), or at least that they display behaviors which would make it appear as though they have them. While this study did not aim to assess the extent, robustness, and human-likeness of the theory-of-mind and world models that LLMs may or may not possess, the performance of LLMs in this study provides some initial evidence for such capabilities, which future studies can assess and quantify.

As noted previously, LLMs displayed different biases than humans in the mirative polarity selection task. Humans showed a small but non-significant preference towards selecting mirative markers (over anti-mirative markers). On the other hand, Meta-Llama models showed a significant mirative preference, GPT-4 showed a significant anti-mirative preference, and GPT-4o showed no preference. Bayesian inference can explain the mirative bias (in humans and Meta-Llama models): when the stimulus lacks sufficient information to determine its label, prior knowledge must be used instead, and the base frequency of “surprisingly” is substantially higher than “unsurprisingly” in natural language. Another source of bias observed in LLMs (but not in humans) was the presentation order of choices, in line with previous research^53,54^. Whether considering all participants or only the subset of participants who completed the task under both choice orders (within-subject analysis), no significant order bias was found in humans. In contrast, the Meta-Llama models and GPT-4o displayed a significant preference towards the first choice presented, and GPT-4 displayed a significant preference towards the last choice presented. The combination of a last-choice bias and an anti-mirative bias renders GPT-4 as particularly un-human-like in this task. The biases exhibited by LLMs suggest that although LLMs achieve comparable accuracy to humans, they arrive at their response through a different underlying process.

### Limitations

This study has several limitations. First, there are several known drawbacks of using online questionnaires administered through the internet. Although participants were instructed to dedicate 20 minutes of their time and complete the tasks in an environment free from distractions, there is no way to verify that this occurred. Nonetheless, participants’ responses in the sentence-completion task showed no indication of randomly typing words or misreading the sentence fragments, and participants’ performance on the mirative-polarity selection task was high. This suggests that, overall, participants were engaged with the tasks.

Another limitation is that, in the mirative-polarity selection task, we assumed that each stimulus contains an expectation violation (or confirmation, for the anti-mirative label). This assumption is somewhat justified since the stimuli were naturally produced, but there is no way to independently verify this. Moreover, since the stimuli were sampled from the internet, the identity of their authors (and relevant information such as whether they are fluent English speakers) is unknown, although we did select stimuli that appeared well-formed and contained no clear misspellings. We take top human performance (90% accuracy) as a tentative approximation that 90% of the stimuli were discernible in nature and thus contained an expectation violation or confirmation.

Finally, in this study, we treated the presence of a mirative marker as a sincere indicator of a violation of expectations. However, in natural text, sarcasm allows speakers to use markers to convey their opposite meaning^64^. While we believe this usage is rare, we cannot discount the possibility of human participants or LLMs treating the mirative marker (or the anti-mirative marker) as sarcastic, which may add noise to our measurements.

## Conclusions

This study researched the sensitivity of humans and LLMs to mirativity, the explicit marking of expectation violations. The two parts of this study can be viewed as investigating the two sides of discourse: the receiver of a textual message and the transmitter of the message. In Part 1, we examined expectation modification after a mirative marker is encountered, which is analogous to the receiving end of discourse. In Part 2, we examined the selection between mirative and anti-mirative connectors for a given sentence pair, which is analogous to producing and transmitting a message, and signaling the level of unexpectedness of a message (in context) to the receiver. We found that both humans and LLMs use mirative markers as cues for calibrating their subsequent expectations during sentence processing, and both humans and LLMs accurately identify when mirative markers are appropriate, albeit with different biases. Together, these findings highlight the active role of expectation management processes during sentence comprehension, and demonstrate how mirative markers modulate the process of communicating unexpected information.

## Methods

### Part 1: Completing sentence fragments with mirative markers

#### Participants

A total of 40 native English speakers with no diagnosed dyslexia or learning disabilities participated in this part of the study by completing an online questionnaire. Participants were recruited through Facebook posts and word of mouth. All participants completed high-school in an English-speaking country. All participants electronically signed an informed consent form and were compensated for their participation.

#### Stimuli

For our main analysis we created 30 novel sentence fragments. The sentence fragments were 4-10 words long, all contained a subject and a verb, and all terminated just before the object appears. The sentence fragments were constructed such that there was at least one appropriate object that could plausibly continue the sentence, although in many cases there was more than one plausible continuation. Each sentence fragment appeared in two conditions: the *neutral* condition where the sentence fragment was preceded by a neutral marker (“yesterday”), and the *mirative* condition where the sentence fragment was preceded by a mirative marker (“surprisingly”).

### Procedure

#### Human

Participants performed an online sentence-completion (cloze) task. They were instructed to type a plausible word that would continue each sentence fragment. Participants were presented with each sentence fragment once: 15 sentences in the neutral condition and 15 in the mirative condition, presented in a pseudo-random order (no more than 3 consecutive trials of the same condition). The assignment of sentence fragments to conditions was counterbalanced between participants. Due to an unfortunate set-up error, one sentence fragment was presented in the mirative condition only, and was excluded from analyses. Responses were recorded while correcting for any typos (e.g., shampane was replaced with the correct spelling, champagne).

#### LLM

The following LLMs were included: Llama-3-8B/70B, Llama-3.1-8B/70B, and Llama-3.2-1B/3B (base versions)^65^. These LLMs were chosen because they are open source, allowing us to calculate the probability of the top token and the overall entropy. The LLMs were accessed with permission through the *Hugging Face transformers* library^66^. LLMs were presented with each sentence fragment twice, once per condition. For each sentence fragment, the full next token probability distribution was recorded.

The LLM was re-initialized between trials to ensure that no conversational history or prior context influenced the responses.

### Analysis

#### Human data

We identified the most popular response under the neutral condition (‘top response’), then calculated its response probability under the neutral condition and the mirative condition. We also calculated the entropy of the response distribution per sentence fragment, per condition. We then tested if the inclusion of a mirative marker modifies the probability of the ‘top response’ and the next word entropy using two-tailed paired t-tests. Probabilities were logit-transformed prior to statistical testing^67^.

#### LLM data

For each sentence fragment, we calculated the next token entropy (calculated over the entire vocabulary) for the neutral and mirative conditions. We also recorded the top 10K tokens under the neutral condition and their probability in the neutral condition and the mirative condition. Similarly to the human case, we then tested if the inclusion of a mirative marker modifies the probability of the ‘top response’ and the next token entropy, using two-tailed paired t-tests. Probabilities were logit-transformed^67^. We also inspected the mean next token probability per marker, calculated across all sentence fragments, for each of the top 10K tokens produced in the neutral condition. Data points were smoothed using locally estimated scatterplot smoothing (LOESS) with span=10.

#### Follow-up testing and analyses (LLM only)

We conducted a follow-up analysis to investigate the scope of the expectation-modification effect of mirative markers and the factors that modulate it. To this end, we used Llama-3.1-8B, the largest open-source LLM we tested, which has shown a significant increase in entropy comparable to human participants. **Stimuli:** We selected 30 passages from the Provo corpus^49^. We selected passages that contained a minimal number of punctuation marks and non-English characters, and no connectors (such as “however”, “for example”) or other markers as the first word of the passage (which would have prevented us from adding a mirative marker in that position). We also made sure that the passage was not explicitly about expectations or surprises. Passages were 2-3 sentences long (39-62 words, mean sentence length 21.35 words). Passages were then converted into all possible sentence fragments, 3 words and longer, starting at the beginning of the passage (e.g., “All that the”, “All that the brain”, “All that the brain has”, “All that the brain has to”, and so on…). Overall, 1,365 sentence fragments were generated from the original 30 passages. Each sentence fragment appeared in two conditions: the neutral condition, where sentence fragments began as they were found in the corpus, or the mirative condition, where sentence fragments began with a mirative marker (“surprisingly”) that was added to the passage. **Procedure:** The procedure was identical to the main analysis. The LLM was presented with each sentence fragment twice, once per condition. The LLM was re-initialized between trials to ensure that no conversational history or prior context influenced the responses. **Analyses:** For each sentence fragment, we calculated the difference in the next token entropy between the mirative and neutral conditions (positive values indicate higher entropy in the mirative condition). To assess the scope of the mirative marker’s effect on expectations, we used two separate methods. First, we fit a linear mixed-effects model to the differences in entropy, with a fixed effect of sentence number within the passage and random intercepts per passage. Second, we fit a LOESS regression (with span=20), predicting the difference in entropy based on the distance (number of words) from the beginning of the passage. These methods allowed us to locate a “window of altered expectations”, defined as the location and range of the increased entropy following a mirative marker. The size of the window could be measured in number of sentences or in number of words. Finally, we further assessed two additional factors which may influence the entropy: the POS of the ‘top response’ (and whether it is a content or a function word), and the level of contextual constraint per sentence fragment (i.e., to what extent does the sentence fragment limits the upcoming tokens). Taking only the data within the “window of altered expectations”, we then fit a linear mixed-effect model to the differences in entropy and tested for the significance of the fixed effects. The model included the predicted token’s syntactic POS (obtained using an automatic POS tagger^68^), the entropy in the neutral condition (which is taken as a measure for the level of contextual constraint), a control variable of a cubic B-spline for the word number within the passage, and random intercepts per passage. A post-hoc test was performed, using estimated marginal means, to compare content vs. function words. All regressions were conducted using the *lme4* package^69^ in R^70^.

### Part 2: Identifying expectation violations

#### Participants

A total of 63 native English speakers with no diagnosed dyslexia or learning disability participated in this part of the study by responding to an online questionnaire. Participants were recruited through Facebook posts and word of mouth. All participants completed high-school in an English-speaking country. Two different versions of the online questionnaire were presented, with a different order of the two presented choices. A subset of 22 participants completed both versions of the questionnaire. One participant was excluded from this study due to not completing the questionnaire. All participants electronically signed an informed consent form and were compensated for their participation.

#### Stimuli

For our main analysis, we selected 1000 sentence pairs from the Discovery discourse marker dataset^50^, 500 per label: for the *mirative* label, sentence pairs were connected with a mirative marker (“surprisingly”), while for the *anti-mirative* label, sentence pairs were connected with an anti-mirative marker (“unsurprisingly”). The sentences were originally taken from a corpus of natural text, and the mirative or anti-mirative connector was embedded between the sentences in the original text. Thus, the connector that actually appeared in the original text constituted the correct label for that pair. A reason for using sentence pairs as the stimuli in this task is that pair structure lends itself to setting up an expectation in the first sentence, indicating a violation of that expectation with a mirative marker connector, and conveying the violation in the second sentence. Sentence pairs were selected such that they contained minimal punctuation marks or non-English characters, the sentences were well-formed, there were no clear misspellings, and there were no sentences that clearly came from “non-communicative” texts such as movie scripts or website menus. Stimulus selection was performed blindly, without knowing the correct label (mirative or anti-mirative), in order to avoid any potential bias.

### Procedure

#### Human

To estimate human level performance on this task, two lists of 30 randomly sampled sentence pairs (15 per label in each list) were presented to human participants through an online questionnaire. Sentence pairs were presented in a 2-alternative forced choice paradigm as follows: the first sentence in the pair was presented, and underneath it appeared two possible continuations: the second sentence in the pair preceded by a mirative marker or by an anti-mirative marker (e.g., “You will only see locals at this restaurant. A) Surprisingly, they have menus in English. B) Unsurprisingly, they have menus in English.”) Participants were instructed to select the most natural continuation out of the two choices (A or B). There were two possible versions of the questionnaire, one where the choice order presented was mirative marker first, and another where the choice order presented was anti-mirative marker first. A subset of participants responded to both questionnaire versions, presented with different sentence pairs in each session, with a break of at least a week between sessions.

#### LLM

The following LLMs were included: Llama-3-8B-Instruct, Llama-3.1-8B-Instruct, Llama-3.2-1B/3B-Instruct^65^, GPT-3.5, GPT-4/4-0613^71^, and GPT-4o/4o-mini^72^ (“chat” versions). The Meta-Llama series of LLMs was accessed with permission through the *Hugging Face transformers* library^66^. The chatGPT series of LLMs was accessed using the OpenAI API. LLMs were presented with each stimulus as a prompt and their text response was recorded. Each prompt began with the instructions: “Please select the most natural continuation of the text from the multiple-choice list. Respond with your choice only.” The instructions were followed by the first sentence and two possible choices, which were the second sentence of the pair either connected with a mirative or anti-mirative marker, exactly like the presentation for the human participants. We looked for LLMs’ responses to contain the second sentence with their chosen connector. Responses were set to be deterministic and we always considered the top token returned as the LLM’s response. For the chatGPT series of LLMs, which are closed-source and cannot be made to respond deterministically, the text response did not always match the top token which was provided separately in the output. In those cases, we coded the LLM response as the top token provided, such that if it was the beginning of the mirative marker (*Sur/sur/ Sur/ sur*) the response was coded as mirative and if it was the beginning of the anti-mirative marker (*Un/un/ Un/ un*) the response was coded as anti-mirative, taking into account different capitalizations and possible inclusions of preceding spaces. If the response returned by the LLM did not begin with any of the markers, it was coded as an invalid response and excluded from later accuracy calculations. Each LLM was presented with all the sentence pairs twice, once where the choice order presented was mirative marker first, and another where the choice order presented was anti-mirative marker first.

### Analysis

#### Human data

Using the participants’ responses, we calculated the accuracy and the response rate per label for each participant, and from that we calculated the mean and standard error (SE) across all participants. We also looked at the accuracy of the top performing participants as an additional benchmark to compare LLM performance against. Agreement between participants was calculated using Gwet’s AC1 metric^73^.

#### LLM data

Accuracy and response rate per label were calculated. An additional stricter measure of accuracy, which we called j*oint order accuracy*, was calculated by considering the LLM response as correct only if it selected the correct label for a given sentence pair under both presentations with different order of choices. We then compared LLM performance with the mean human performance and the top human performance.

### Follow up testing and analyses (LLM only)

#### Data discriminability

To ensure that the high performance of the LLMs was derived from some ability to identify expectation violations and not by relying on other heuristics, we measured the discriminability of the data using several simple variables: the length of the first sentence or the second sentence (measured in tokens using the Llama-3 tokenizer), the perplexity and the maximum-surprisal (max-surprisal) of the second sentence given the first sentence without any connector (measured using Llama-3-8B base version), the cosine similarity between the first and second sentence (measured using OpenAI’s embeddings model text-embedding-3-small), whether the first sentence was a question or not, and whether the first or second sentence contained any negativity markings (i.e., whether the sentence pair includes any of the following words: *no/not/_n’t/none/nothing/never/neither/cannot)*.

Discriminability was measured using the receiver operating characteristic (ROC) and the area under the curve (AUC) was calculated for each variable.

#### Manipulated stimuli

An additional test, using manipulated stimuli, was conducted on the best performing LLMs (Llama-3-8B-Instruct, Llama-3.1-8B-Instruct, chatGPT-4, chatGPT-4o). The manipulated stimuli included: (a) only the first sentence of each pair, (b) only the second sentence of each pair, and (c) sentence pairs generated by shuffling randomly the first and second sentences within a given label. Once again, each LLM was presented with all the sentence pairs twice: once with the choice of a mirative marker presented first, and another with the choice of an anti-mirative marker presented first.

#### Biases

Beyond comparing human and LLM task performance, we also compared their respective biases in task performance. Two sources of bias were considered: the ‘mirative bias’, the propensity to select a mirative marker over the alternative, and the ‘order bias’, the propensity to select the last choice presented. We estimated the biases using a logistic mixed-effects regression, using the mirative response rate as the dependent variable, an intercept and choice order as the independent variables, and another random intercept per stimulus. For human participants, we also included a random intercept per participant. The intercept and order coefficients extracted from the regression model were interpreted as the log odds-ratio to select the mirative marker over the anti-mirative marker and the log odds-ratio to select the second choice presented over the first choice presented, respectively. A more conservative estimate for the human bias was calculated in the same manner but using only the 22 participants who completed both versions of the questionnaire. All regressions were conducted using the *lme4* package^69^ in R^70^.

## Supporting information

Supplementary analysis

## Acknowledgements

This study was conducted as part of Benjamin Menashe’s doctoral dissertation, under the supervision of Prof. Michal Ben-Shachar at the Gonda Multidisciplinary Brain Research Center, Bar-Ilan University. This project was supported by the Data Science Institute at Bar-Ilan University. B.M. is supported in part by the President’s scholarship of BIU. We thank Prof. Yael Greenberg and Prof. Yoav Goldberg for many thoughtful discussions.

## Author contributions

Benjamin Menashe ran the experiments, analyzed the data, and wrote the manuscript. Austin Drake developed stimuli and helped review the manuscript. Michal Ben-Shachar supervised this study.

## Competing interests

The authors declare no competing interests.

## Data availability statement (mandatory)

The datasets generated and analyzed during the current study will be made available in an open-source public repository [LINK].

## Notes

### Competing Interest Statement

The authors have declared no competing interest.

### Summary of Updates

fixed minor typos; few sentences edited for clarity.

