## Supplementary Table for "Expectation management in humans and LLMs"

### Supplementary Tables

#### Table S1. Handcrafted stimuli

| model | comparison | difference | $t_{29}$ | p-value | |
|---|---|---|---|---|---|
| Human | entropy | 0.628 ± 0.11 | 5.58 | <.001 | *** |
| | p(top response) | −1.149 ± 0.17 | −6.85 | <.001 | *** |
| Llama-3-8B | entropy | 0.523 ± 0.17 | 3.01 | .005 | ** |
| | p(top token) | −0.656 ± 0.13 | −5.02 | <.001 | *** |
| Llama-3-70B | entropy | 0.098 ± 0.19 | 0.53 | .601 | |
| | p(top token) | −0.724 ± 0.19 | −3.74 | .001 | ** |
| Llama-3.1-8B | entropy | 0.648 ± 0.19 | 3.45 | .002 | ** |
| | p(top token) | −0.695 ± 0.13 | −5.39 | <.001 | *** |
| Llama-3.1-70B | entropy | 0.226 + 0.20 | 1.11 | .277 | |
| | p(top token) | −0.860 + 0.20 | −4.28 | <.001 | *** |
| Llama-3.2-1B | entropy | 0.216 ± 0.13 | 1.65 | .109 | |
| | p(top token) | −0.466 ± 0.17 | −2.73 | .011 | * |
| Llama-3.2-3B | entropy | 0.483 ± 0.15 | 3.24 | .003 | ** |
| | p(top token) | −0.707 ± 0.15 | −4.77 | <.001 | *** |

Mean ± SE where applicable.
Probabilities are compared on the logit scale.

**Table S1. Increased entropy and reduced probability in all LLMs tested.** The presence of a mirative marker in the beginning of the sentence fragment significantly decreased the probability of the most-expected response in all six LLMs tested. It also decreased the entropy, although the decrease in entropy was significant in only three of the six LLMs tested. * p<.05, ** p<.01, *** p<.001.

**Table S2. Summary of linear mixed-effects models**

| model | data | variable | (level) | marginal mean | $t_{df}$ | p-value |
|---|---|---|---|---|---|---|
| model 1 | all | sentence number | sentence 1 | 0.176 ± 0.03 | $5.16_{38.4}$ | <.001 *** |
| | | | sentence 2 | 0.128 ± 0.03 | $3.74_{39.4}$ | <.001 *** |
| | | | sentence 3 | 0.050 ± 0.04 | $1.18_{82.4}$ | .243 |
| model 2 | mirativity window | word POS | adjective | 0.311 ± 0.08 | $3.99_{470.7}$ | <.001 *** |
| | | | adverb | 0.222 ± 0.09 | $2.49_{586.1}$ | .013 * |
| | | | determiner | 0.223 ± 0.06 | $4.04_{173.8}$ | <.001 *** |
| | | | noun | 0.266 ± 0.05 | $5.43_{114.9}$ | <.001 *** |
| | | | preposition | 0.224 ± 0.05 | $4.12_{167.0}$ | <.001 *** |
| | | | pronoun | 0.240 ± 0.07 | $3.42_{372.6}$ | <.001 *** |
| | | | verb | 0.176 ± 0.06 | $3.07_{200.3}$ | .002 ** |
| | | neutral condition entropy | | −0.002 ± 0.07 | $-0.29_{849.7}$ | .774 |
| | | distance from marker (B-Spline) | 5 | 0.042 ± 0.08 | $0.54_{509.0}$ | .592 |
| | | | 10 | 0.160 ± 0.05 | $3.46_{96.9}$ | <.001 *** |
| | | | 15 | 0.213 ± 0.05 | $4.66_{91.3}$ | <.001 *** |
| | | | 20 | 0.242 ± 0.05 | $5.32_{86.7}$ | <.001 *** |
| | | | 25 | 0.193 ± 0.04 | $4.49_{68.9}$ | <.001 *** |
| | | | 30 | 0.122 ± 0.05 | $2.60_{98.0}$ | .011 * |
| | | | 35 | 0.098 ± 0.05 | $1.88_{137.8}$ | .061 |
| | | | 40 | 0.096 ± 0.09 | $1.13_{562.9}$ | .258 |

Mean ± SE where applicable. DF computed using Satterthwaite's method.

**Table S2. Summary of linear mixed-effects models. Model 1)** We fit a linear mixed-effects model on all words from the 30 texts. Our dependent variable was the difference in entropy between the mirative condition and the neutral condition. Entropy was estimated using meta-llama-3.1-8B. The linear mixed-effects model included the sentence number in the text and a random intercept per text. Reported are the estimated marginal means per sentence number. **Model 2)** We fit a linear mixed-effects model using the words inside the mirativity window (within 2 sentences and a distance of 5-40 words from the mirative marker). Our dependent variable was the difference in entropy between the mirative condition and the neutral condition. The model included the predicted token's POS, the entropy in the neutral condition, a control variable of a cubic B-spline for the word's distance from the mirative marker, and a random intercept per text. Reported are the estimated marginal means per level for the word POS coefficients, the estimated marginal trend for the neutral condition entropy, and the estimated marginal mean for various distances from the mirative marker. All estimates per coefficient are reported after averaging over the levels or trends of the other coefficients.

**Table S3. Mirative polarity selection – summary of responses**

| model | mean accuracy | joint order accuracy | invalid responses | first choice presented | accuracy | mirative accuracy | anti-mirative accuracy | mirative response rate | anti-mirative response rate | mirative bias | order bias |
|---|---|---|---|---|---|---|---|---|---|---|---|
| Human | **75.5%** ± 1% | | | mirative | **74.9%** ± 2% | 83.5% ± 2% | 66.3% ± 4% | 58.6% ± 2% | 41.4% ± 2% | 0.51 ± 0.3 | −0.02 ± 0.1 |
| | | | | anti-mirative | **76.1%** ± 2% | 81.6% ± 2% | 70.5% ± 3% | 55.6% ± 2% | 44.4% ± 2% | | |
| Llama-3-8B-Instruct | **65.2%** | 40.4% | 0.7% | mirative | **63.9%** | 87.7% | 40.2% | 73.7% | 26.3% | 0.60 ± 0.1 | −0.87 ± 0.1 |
| | | | | anti-mirative | **66.4%** | 70.4% | 62.4% | 54.0% | 46.0% | | |
| Llama-3.1-8B-Instruct | **67.9%** | 41.5% | 1.8% | mirative | **65.7%** | 88.0% | 43.5% | 72.3% | 27.7% | 0.31 ± 0.1 | −1.43 ± 0.1 |
| | | | | anti-mirative | **70.0%** | 60.8% | 79.3% | 40.9% | 59.1% | | |
| Llama-3.2-1B-Instruct | **47.4%** | 24.6% | 35.1% | mirative | **48.2%** | 0.5% | 99.7% | 0.4% | 99.6% | −14.4 ± 1.6 | 6.04 ± 1.6 |
| | | | | anti-mirative | **46.4%** | 1.6% | 97.8% | 1.8% | 98.2% | | |
| Llama-3.2-3B-Instruct | **52.1%** | 6.3% | 0.5% | mirative | **54.3%** | 12.6% | 96.0% | 8.3% | 91.7% | 1.56 ± 0.3 | 7.92 ± 0.5 |
| | | | | anti-mirative | **49.9%** | 99.6% | 0.4% | 99.6% | 0.4% | | |
| GPT-3.5-turbo-0125 | **61.3%** | 26.4% | 0.5% | mirative | **55.7%** | 96.4% | 14.6% | 90.9% | 9.1% | 0.64 ± 0.1 | −3.44 ± 0.2 |
| | | | | anti-mirative | **66.8%** | 42.8% | 91.0% | 26.0% | 74.0% | | |
| GPT-4-0613 | **76.9%** | 62.8% | 0.2% | mirative | **72.4%** | 49.3% | 95.4% | 26.9% | 73.1% | −1.06 ± 0.1 | 1.62 ± 0.2 |
| | | | | anti-mirative | **81.3%** | 78.6% | 84.0% | 47.2% | 52.8% | | |
| GPT-4-turbo-2024-04-09 | **78.5%** | 61.7% | 0.0% | mirative | **72.2%** | 46.8% | 97.6% | 24.6% | 75.4% | −0.88 ± 0.1 | 2.59 ± 0.2 |
| | | | | anti-mirative | **84.7%** | 90.2% | 79.2% | 55.5% | 44.5% | | |
| GPT-4o-mini-2024-07-18 | **68.7%** | 48.3% | 0.3% | mirative | **61.6%** | 97.4% | 25.7% | 85.9% | 14.1% | 1.20 ± 0.1 | −2.23 ± 0.2 |
| | | | | anti-mirative | **75.9%** | 77.5% | 74.3% | 51.6% | 48.4% | | |
| GPT-4o-2024-08-06 | **84.8%** | 77.1% | 0.0% | mirative | **84.3%** | 87.0% | 81.6% | 52.7% | 47.3% | 0.12 ± 0.1 | −0.32 ± 0.1 |
| | | | | anti-mirative | **85.2%** | 84.6% | 85.8% | 49.4% | 50.6% | | |

Mean ± SE where applicable.

**Table S3. Mirative polarity selection – summary of responses.** Mirative polarity selection is a novel task where participants and LLMs are presented with a sentence pair and asked to select whether it was connected with a mirative marker (“surprisingly”) or an anti-mirative marker (“unsurprisingly”). We used 1000 sentence pairs (500 per marker) from the Discovery dataset (Sileo et al., 2019). Human participants were presented with 60 randomly selected stimuli (30 per marker). Human participants and models were presented with stimuli where the choices were presented either with the mirative marker first or with the anti-mirative marker first. “Mean accuracy” indicates the mean accuracy level over both choice orders. “Joint order accuracy” assigns a “correct” response only when the model responded correctly for both choice orders. “Invalid responses” are cases where the model did not respond with selecting a mirative or an anti-mirative marker (e.g., “I cannot select an answer”). Accuracy level and response rates are further reported by choice order presented, separately for miratives and anti-miratives, in order to highlight the mirativity bias and order bias LLMs displayed. Mirativity bias and order bias were quantitively assessed by fitting logistic mixed-effects regression models for each LLM, using the mirative response rate as the dependent variable, an intercept and choice order as the independent variables, and a random intercept per stimulus. Similarly, for the human participants we fit a logistic mixed effects regression model with the same specification but with the addition of random intercepts per participant.

**Table S4. Manipulated stimuli**

| condition | model | mean accuracy | joint order accuracy | invalid responses |
|---|---|---|---|---|
| original | Human | **75.5%** ± 1% | | |
| | Llama-3-8B | **65.2%** | 40.4% | 0.7% |
| | Llama-3.1-8B | **67.9%** | 41.5% | 1.8% |
| | gpt-4-turbo-2024-04-09 | **78.5%** | 61.7% | 0.0% |
| | gpt-4o-2024-08-06 | **84.8%** | 77.1% | 0.0% |
| sentence 1 only | Llama-3-8B | **60.2%** | 40.5% | 0.2% |
| | Llama-3.1-8B | **58.2%** | 44.1% | 0.0% |
| | gpt-4-turbo-2024-04-09 | **62.6%** | 42.2% | 0.0% |
| | gpt-4o-2024-08-06 | **62.2%** | 45.9% | 0.0% |
| sentence 2 only | Llama-3-8B | **52.0%** | 13.4% | 3.8% |
| | Llama-3.1-8B | **52.3%** | 7.4% | 29.2% |
| | gpt-4-turbo-2024-04-09 | **66.2%** | 39.6% | 0.1% |
| | gpt-4o-2024-08-06 | **69.1%** | 50.6% | 0.1% |
| shuffled | Llama-3-8B | **60.8%** | 28.0% | 9.0% |
| | Llama-3.1-8B | **57.7%** | 22.6% | 1.6% |
| | gpt-4-turbo-2024-04-09 | **62.2%** | 31.7% | 5.6% |
| | gpt-4o-2024-08-06 | **63.3%** | 25.6% | 19.7% |

Mean ± SE where applicable.

**Table S4. Mirative polarity selection – manipulated stimuli.** Summary of the model responses for the manipulated stimuli conditions: original (no manipulation), using the first sentence only, using the second sentence only, and randomly shuffling the first and second sentences of different sentence pairs within the same marker. Results are averaged across both choice orders. Mean accuracy is the mean accuracy over both choice orders. Joint order accuracy is the accuracy only if the model responded correctly for both choice orders. Invalid responses are the percent of response where the model did not respond with selecting a mirative or an anti-mirative marker.

#### Table S5. Example sentence pairs

| sentence pair | label | human accuracy |
|---|---|---|
| One reason for this could be that they're happy to see me go. **Unsurprisingly,** I don't think this is true. | anti-mirative | 34.9% |
| For those of you wanting to see some Hindi cinema, this is about as far from Hindi cinema as a Hindi film can get. **Unsurprisingly**, there are no songs. | anti-mirative | 37.2% |
| I also wanted a crash course on world culture, and some supplementary materials I could use to help diversify my lesson plans. **Surprisingly**, I received almost none of this. | mirative | 50.0% |
| In an awkward moment, Price asked the jury to raise their hands "if they love Apple". **Unsurprisingly**, the jury did nothing. | anti-mirative | 52.4% |
| The whole landlord and business class was robbed of its social power and status across much of Asia and Europe. **Unsurprisingly,** this generated much bitterness. | anti-mirative | 90.5% |
| It's actually quite interesting to see what these aliens actually have in means of reproductive systems. **Surprisingly**, things look similar enough to our own. | mirative | 97.6% |
| Picked this game up last Saturday on the Shop Channel not knowing what I'd get myself into. **Surprisingly**, I found this to be great fun. | mirative | 97.7% |

**Table S5. Mirative polarity selection - examples.** Examples of sentence pairs presented to human participants in the mirative polarity selection task. Examples include some sentence pairs with low accuracy, chance level accuracy, and high accuracy.
