## Supplementary analysis for "Expectation management in humans and LLMs"

In a supplementary analysis, we wanted to see if the success of LLMs on the mirative polarity selection task arises because of some "common-sense reasoning" ability associating the mirative marker with an expectation violation. To do so, we tested the impact of including a mirative marker in two other tasks: a common-sense reasoning task and a vocabulary task.

For the common-sense reasoning task, we selected 100 false factual statements from the com2sense dataset[41] (see example in Supplementary Fig. S1, panel A1). Importantly, all statements presented false information as factual, thus mimicking a violation of expectations. We added a neutral marker ("additionally") or a mirative marker ("surprisingly") to the beginning of each statement. LLMs (Llama-3-8B-Instruct, Llama-3.1-8B-Instruct, chatGPT-4, chatGPT-4o) were instructed to respond to each statement as "true" or "false". Since all statements were false, lower accuracy on the task indicates that more statements were perceived as true. We also recorded the probability of the response "false" produced by the Llama models, for all stimuli where the models responded correctly.

For the vocabulary task, 499 vocabulary question were taken with permission from a college-prep book (Chesla, 2007; (see example in Supplementary Fig. S1, panel B1). Vocabulary questions included a sentence with a missing word presented as blank, and a multiple choice of five low frequency words, only one of which fits semantically in the sentence. We added a neutral marker ("additionally") or a mirative marker ("surprisingly") to the beginning of each question. LLMs (Llama-3-8B-Instruct, Llama-3.1-8B-Instruct, chatGPT-4, chatGPT-4o) were instructed to respond with the correct vocabulary word for each sentence. We also recorded the probability of the correct word from the Llama models, for all stimuli where the model responded correctly. If the correct word was tokenized into more than one token, the probability of that word was calculated by multiplying the probability for each token, given all previous tokens.

We then calculated the difference in logit-transformed probability between the neutral condition and mirative condition (equivalent to the log odds-ratio) for each stimulus, in both tasks. A more negative log odds-ratio indicates that the addition of a mirative marker lowers the probability assigned to the correct response, as compared to the neutral marker. We tested if the log odds-ratio is significantly different in the common-sense reasoning task as compared to the vocabulary task. After finding that the distributions deviate significantly from normality (Shapiro-Wilk test, all p<.01), we conducted a two-tailed Mann-Whitney U test, separately for each model.

Importantly, the tasks were chosen such that sensitivity to mirativity should affect performance on the common-sense reasoning task but not on the vocabulary task. Thus, we predicted that the presence of a mirative marker will affect LLMs in the common-sense reasoning task more than the vocabulary task.

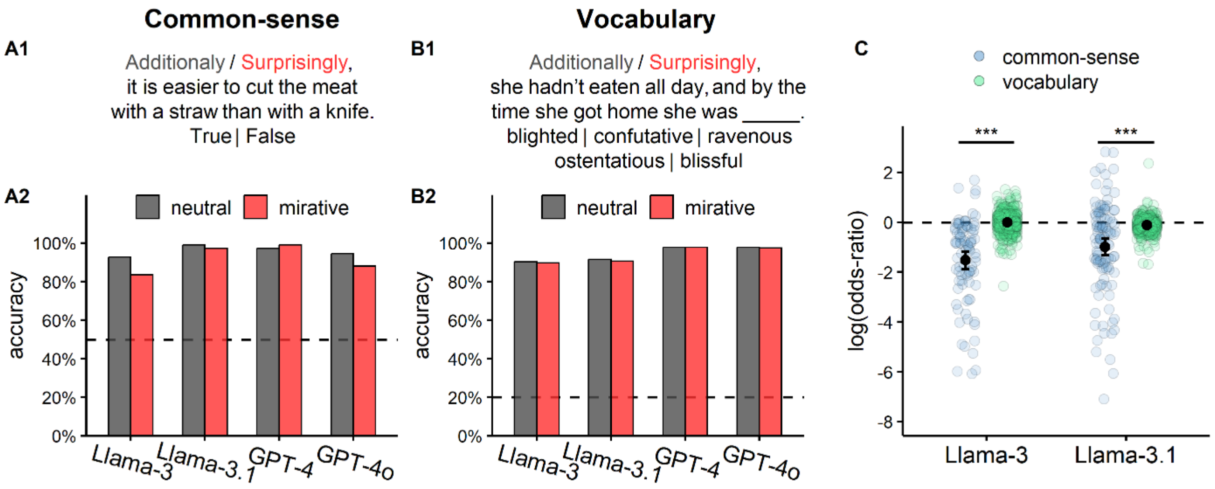


**Supplementary Figure S1. Mirative markers affect common-sense reasoning more than vocabulary. A)** We tested LLMs on a common-sense reasoning task. LLMs were instructed to respond True/False to prompts containing 100 false common sense reasoning questions taken from the Com2Sense dataset (Singh et al., 2021). Questions began either with a neutral marker ("additionally") or a mirative marker ("surprisingly"). **A1)** example stimulus. The correct answer is false. **A2)** Model accuracy per condition. **B)** We tested LLMs on a vocabulary task. LLMs were instructed to respond to prompts containing 499 multiple-choice vocabulary questions (Chesla, 2007). Questions began either with a neutral marker ("additionally") or a mirative marker ("surprisingly"). **B1)** Example stimulus. The correct answer is "ravenous". **B2)** Model accuracy per condition. **C)** Log odds-ratios of the model probabilities for the correct responses, per task. Negative values indicate a lower probability of the correct response on the mirative condition as compared to the neutral conditions. *** p<.001.

Results show that the presence of a mirative marker slightly decreased the accuracy on the common-sense reasoning task in 3 of the 4 LLMs tested (percent drop in accuracy: Llama-3-8B-Instruct: 9.1%. Llama-3.1-8B-Instruct: 1.8%. GPT-4-turbo-2024-04-09: -1.8%. GPT-4o-2024-08-06: 6.4%), indicating that the mirative marker increases the tendency to classify false statements as true. This is likely due to the fact that false statements about the world tend to violate expectations about the world, and thus the presence of a mirative marker can signal that although the statement is violating expectations, it still may be true. However, the presence of a mirative marker did not affect performance on the vocabulary task (percent drop in accuracy: Llama-3-8B-Instruct: 0.4%. Llama-3.1-8B-Instruct: 1.0%. GPT-4-turbo-2024-04-09: 0%. GPT-4o-2024-08-06: 0.4%; See Supplementary Fig. S1, panels A2 and B2, for LLM performance by marker). We further examined the probability that models provided for the correct response. The presence of a mirative marker decreased the probability for the correct response in the common-sense reasoning task significantly more than in the vocabulary task (mean log odds-ratio: Llama-3-8B-Instruct: common-sense = −1.45, vocabulary = −0.01, $p<.001$; Llama-3.1-8B-Instruct: common-sense = −1.00, vocabulary = −0.12, $p<.001$; See Supplementary Fig. S1C for a comparison of the probability for the correct response by task). This result shows that mirative markers affect LLMs in tasks where common-sense reasoning is involved to a much greater degree than in tasks which involve some other lexical ability. This provides support for the claim that LLMs are sensitive to mirativity and process mirative markers as indicators of expectation violation.
